# Designing Cell-Type-Specific Promoter Sequences Using Conservative Model-Based Optimization

**DOI:** 10.1101/2024.06.23.600232

**Authors:** Aniketh Janardhan Reddy, Xinyang Geng, Michael H. Herschl, Sathvik Kolli, Aviral Kumar, Patrick D. Hsu, Sergey Levine, Nilah M. Ioannidis

## Abstract

Gene therapies have the potential to treat disease by delivering therapeutic genetic cargo to disease-associated cells. One limitation to their widespread use is the lack of short regulatory sequences, or promoters, that differentially induce the expression of delivered genetic cargo in target cells, minimizing side effects in other cell types. Such cell-type-specific promoters are difficult to discover using existing methods, requiring either manual curation or access to large datasets of promoter-driven expression from both targeted and untargeted cells. Model-based optimization (MBO) has emerged as an effective method to design biological sequences in an automated manner, and has recently been used in promoter design methods. However, these methods have only been tested using large training datasets that are expensive to collect, and focus on designing promoters for markedly different cell types, overlooking the complexities associated with designing promoters for closely related cell types that share similar regulatory features. Therefore, we introduce a comprehensive framework for utilizing MBO to design promoters in a data-efficient manner, with an emphasis on discovering promoters for similar cell types. We use conservative objective models (COMs) for MBO and highlight practical considerations such as best practices for improving sequence diversity, getting estimates of model uncertainty, and choosing the optimal set of sequences for experimental validation. Using three relatively similar blood cancer cell lines (Jurkat, K562, and THP1), we show that our approach discovers many novel cell-type-specific promoters after experimentally validating the designed sequences. For K562 cells, in particular, we discover a promoter that has 75.85% higher cell-type-specificity than the best promoter from the initial dataset used to train our models.

## 1. Introduction

Gene therapies treat diseases through the delivery and expression of therapeutic genetic cargo in disease-associated cells and tissues. Expression is controlled by a promoter sequence, a short regulatory DNA sequence (typically up to a few hundred base pairs (bp) long) placed upstream of the coding region. An ideal promoter for gene therapy would differentially induce expression in targeted cells while repressing expression in all other cells to increase effectiveness and reduce side-effects; i.e. it should be cell-type-specific and induce differential expression. However, although there are over 400 types of cells in the human body (Tabula Sapiens Consortium, 2022), very few cell-type-specific promoters are known.

Traditional methods to design cell-type-specific promoters rely heavily on manual curation or involve tiling known cis-regulatory elements (CREs) or transcription factor (TF) binding motifs ((Miao et al., 2000; Selvakumaran et al., 2001; Yun et al., 2008; Nissim et al., 2017; Wu et al., 2019) among others). These methods are difficult to automate and are not guaranteed to work, especially in less studied cell types. Directed evolution can be used to automate promoter design but is expensive to perform, typically requiring many experimental validation steps to continually improve designs, and does not make the best use of available data. Offline model-based optimization (MBO) has recently been used to design promoters and other CREs in a data-driven manner (Linder et al. (2020); Wang et al. (2020); Jores et al. (2021); LaFleur et al. (2022); Gosai et al. (2023) among others). The offline MBO approach involves building machine learning (ML) models of promoter-driven expression (PE) using experimental measurements and then optimizing the promoter sequence using model predictions as surrogates for experimental measurements. Offline MBO has the potential to accelerate promoter discovery by being automated and relatively data-efficient; however, the field is lacking a generalizable MBO-based framework for designing cell-type-specific promoters in a data-efficient manner, while systematically accounting for practical considerations including:

### 1. Data-efficiency

Measuring PE is expensive and time-consuming, necessitating the effective use of available data during design. Most previous work on offline MBO for promoter design has used large PE datasets for model training (e.g. from massively parallel reporter assays, MPRAs) that are only available for a few well-studied cell types, and their modelling strategies have not been evaluated when designing cell-type-specific promoters for less-studied cell types using small PE datasets.

### 2. Minimizing adversarial designs

Using models for optimization, rather than just prediction, presents key additional challenges, since naïvely optimizing designs based on predicted differential expression (DE) can lead to *adversarial* designs that fool a model into outputting desirable values while having undesirable values in reality. Previous work using optimization techniques such as *in silico* directed evolution, gradient ascent, or generative models do not account for such designs.

### 3. Selecting design candidates while accounting for sequence diversity and uncertainty in estimates of a sequence’s goodness

Offline MBO can design many sequences that are predicted to be cell-type-specific by the PE model. However, due to limited budgets and time for wetlab experiments, typically only a subset of the candidates can be validated. Thus, we need a systematic approach to choose promising candidates from a larger set of designs in a principled manner. During selection, it is important to consider both the diversity of the chosen candidates and the uncertainty in model predictions. Choosing diverse candidate designs is important (1) to improve the chances of successfully discovering a cell-type-specific promoter upon experimental validation, since we lack complete information about how a sequence controls PE, and (2) to explore sequence space more effectively during several rounds of design, leading to progressively better models and designs. It is also important to choose candidates that have low uncertainty in model predictions while also balancing their potential goodness. This sequence selection process is crucial for a principled MBO workflow and has been overlooked by previous work.

Here we present a systematic framework and case study for applying offline MBO to the real world problem of promoter design, backed by empirical evidence from wetlab experiments evaluating differential expression in three closely related blood-based cell types. To improve data-efficiency through transfer learning, we follow previous work (Reddy et al., 2024) and pretrain sequence-based deep learning (DL) models of PE using large existing MPRA datasets that measure PE in different contexts, and fine-tune them using a small PE dataset collected from the target cell types. To address adversarial designs, we extend the conservative objective models (COMs) framework (Trabucco et al., 2021) to our setting: while fine-tuning models, we use a loss term that reduces the predicted DE of adversarial designs. These fine-tuned models are then used to design cell-type-specific promoters by performing gradient ascent on the inputs. Upon generating multiple candidate sequences, we propose a selection scheme to choose a diverse yet effective subset of final candidates that uses uncertainty estimates from an independently trained ensemble. We use our workflow to design cell-type-specific promoters for three blood cancer cell lines (Jurkat, K562, and THP1) and present experimental evidence of its effectiveness. Existing studies have designed promoters for markedly different cell types derived from different tissues (Gosai et al., 2023), a simpler problem since such cell types typically have very different regulatory environments with different transcription factors (TFs) expressed. We show that our MBO approach works in a more difficult setting where we aim to design promoters that are only expressed in one of three highly similar blood-based cell lines. Our designs improve upon the cell-type-specificity of sequences from the fine-tuning dataset, **improving 72.12% of sequences in Jurkat cells and 80.48% in K562 cells**. Additionally, **in K562 cells, we identify a promoter with 75.85% higher DE than the best promoter** from the fine-tuning dataset. These results suggest that applying our approach over multiple rounds of modelling, de-sign, and experimental validation will enable the discovery of promoters with even higher cell-type-specificity, by shifting the training distribution towards higher DE sequences at each iteration.

## 2. Related work

### 2.1. Promoter design methods

Promoters have traditionally been designed using heuristics and manual curation. For example, Selvakumaran et al. (2001) and Yun et al. (2008) found cell-type-specific promoters for ovarian and breast cancer cells, respectively, by identifying genes that are differentially expressed in these cells and showing that their promoters exhibit cell-type-specificity. Nissim et al. (2017) designed cell-type-specific promoters for ovarian and breast cancer cells by identifying differentially expressed TFs and tiling their binding motifs. However, these studies do not provide a generalizable method to design cell-type-specific promoters for any given cell type. In an attempt to address this issue, recent work has sought to develop more automated and data-driven design methods. Kotopka and Smolke (2020) and Jores et al. (2021) used sequence-based convolutional neural network (CNN) models alongside simple optimization techniques such as *in silico* directed evolution and gradient ascent to design promoters for high expression in yeast and various plant cells, respectively. Wang et al. (2020) generated E. coli promoters by training a generative adversarial network (GAN) on natural E. coli promoters and then using a PE predictor to further filter generated sequences. Gosai et al. (2023) trained a sequence-based CNN model of PE using a large MPRA dataset from three very different cell lines. They used this model with three design algorithms to produce cell-type-specific promoters that they show to be effective and diverse. Here, we build on previous work in three crucial ways. First, we exploit the benefits of pretraining on large existing PE datasets to build accurate models for target cell types in a more data efficient manner compared to previous methods. Second, we leverage COMs (Trabucco et al., 2021), a powerful offline MBO method, to design promoters using these models more effectively, minimizing adversarial designs. Finally, we propose a sequence selection strategy that explicitly accounts for sequence diversity and model uncertainty to choose a subset of sequences for experimental validation from a larger set of design candidates.

### 2.2. Offline MBO for designing biological sequences

Apart from COMs, other offline MBO methods have been proposed for designing biological sequences. While COMs only requires discriminative models, most of the other methods rely on generative models. For example, Brookes et al. (2019) built variational autoencoders (VAEs) that generate desirable designs by iteratively improving the VAE and its designs, guided by a design model. Jain et al. (2022) used GFlowNets for sequence design, another class of generative models that are trained to output designs that span all modes of some design model’s prediction distribution. Linder et al. (2020)’s deep exploration networks (DENs) are generative models that are trained to output diverse yet desirable sequences and again use design model predictions to guide the training process. They experimentally validated their method and showed that alternative polyadenylation (APA) sites designed using DENs were better than those designed using gradient ascent. Here we use DENs as a baseline in our analyses, since it was validated using wetlab experiments.

## 3. Background and motivation

### 3.1. The cell-type-specific promoter design problem

A typical promoter design experiment ((Schlabach et al., 2010; Kotopka and Smolke, 2020; Jores et al., 2021; LaFleur et al., 2022) among others) starts by identifying target cell types and defining an objective such as increasing the absolute or differential expression levels of some gene. Then, an initial set of candidate promoters is prepared by leveraging existing data sources and heuristics, or even using random sequences. To test the efficacy of these promoters, reporter assays that collect PE measurements for a large batch of promoters simultaneously are used (batch sizes range from a few hundred to many thousands or even millions). Then, using these initial PE measurements, better promoters can be designed using various methods such as directed evolution, shuffling motifs enriched in desirable sequences, or offline MBO. The designed promoters then need to be experimentally validated using another reporter assay. If a sufficiently desirable promoter is not discovered, the design and experimental validation steps can be repeated.

Here we focus on designing short cell-type-specific promoters in a data-constrained setting. In this setting, we have a set of target cell types *C*, from which we have a small PE dataset – a few thousand sequences and corresponding PE measurements from all cell types in *C*. Gene delivery mechanisms impose constraints on the length of DNA that can be delivered. Thus, we can often only accommodate promoters that are at most a few hundred base pairs (bp) long. For any target cell type *tc* ∈ *C*, we define the DE of a promoter **x** (i.e. its cell-type-specificity) as:

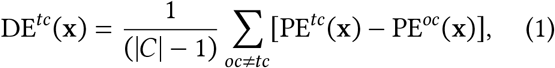

where PE^*c*^(**x**), *c* ∈ *C*, is the experimentally measured expression value induced by **x** in cell type *c*. Our goal is to design sequences that maximize DE^*tc*^ - the average difference in PE between the target cell type *tc* and the other cell types.

### 3.2. Offline MBO

An offline MBO algorithm aims to produce designs that maximize some objective function, using a provided static dataset 𝒟 = {(**x**_*i*_, *y*_*i*_)} of designs **x**_*i*_ and a corresponding measurement of the objective value *y*_*i*_. The algorithm analyzes this dataset and produces an optimized candidate design **x**^*^, which is evaluated against the true objective function. This process often involves learning a proxy objective *f*_*θ*_ mapping designs to corresponding objective values, which we hereafter refer to as the **design model**. Optimization is then performed to find an input which maximizes this learned design model: **x**^*^ = arg max_**x**_ *f*_*θ*_(**x**), using methods such as *in silico* directed evolution, gradient ascent, or by building generative models.

As many offline MBO algorithms are data-efficient, they are suitable for promoter design in data-constrained settings. When several rounds of promoter design are performed, we can use all available PE measurements from previous rounds as the “static” dataset for offline MBO. In our workflow, we propose an additional selection step after running offline MBO that diversifies the final designs if needed, making the pipeline more suitable for multi-round optimization.

## 4. Our MBO workflow for designing promoters in data-constrained settings

In this section, we present our approach to design cell-type-specific promoters that accounts for key practical considerations. Figure 1 summarizes the workflow.

**Figure 1:**
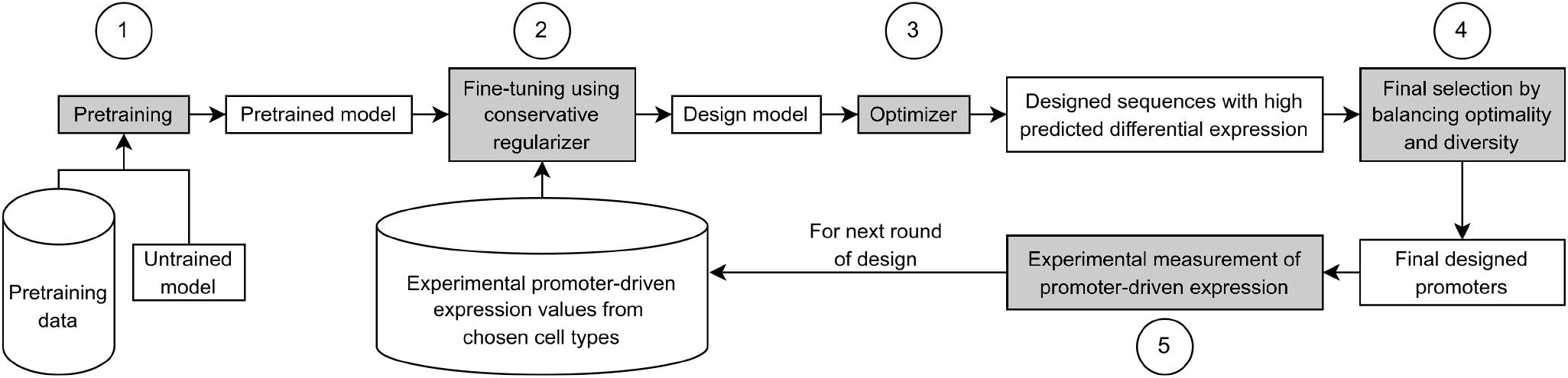
Our workflow for designing cell-type-specific promoters. Five main steps are highlighted in grey: (1) pretrain a base model using existing large genomic datasets; (2) fine-tune the pretrained model using the experimentally measured PE data collected so far, while also using a conservative regularizer; (3) use a gradient ascent-based optimizer to design sequences that have high predicted DE; (4) apply a final sequence selection algorithm that balances optimality and diversity to choose a smaller subset of the designed sequences; (5) experimentally measure the PE of the selected designed sequences. The last four steps can be repeated when running multiple rounds of design iterations.

### 4.1. Building models of promoter-driven expression in a data-efficient manner

As previously mentioned, offline MBO algorithms need a design model that approximates the objective function. When designing cell-type-specific promoters for some target cell type *tc* ∈ *C*, our objective function is DE^*tc*^(**x**) that is in turn computed using all PE^*c*^(**x**), *c* ∈ *C*, values. Therefore, instead of directly modelling DE^*tc*^(**x**), we simultaneously model all PE^*c*^(**x**) values using multi-task learning (MTL). This allows us to use one design model to design cell-type-specific promoters for all *c* ∈ *C*. In this section, we provide guidelines to build these models based on the findings of Reddy et al. (2024), which studies model architectures and transfer learning strategies for modelling PE.

#### Model architecture

Recent work has shown that genomic data can be accurately modelled using sequence-based models – models that take in one-hot encoded DNA sequences as input to predict associated experimental assay measurements such as gene expression, histone modifications, and TF-binding (e.g. (Zhou and Troyanskaya, 2015; Agarwal and Shendure, 2020; Avsec et al., 2021)). Reddy et al. (2024) find that a model consisting of convolutional layers followed by transformer layers (MTLucifer) outperforms other architectures when used to model PE without any transfer learning.

#### Transfer learning to boost accuracy

Effective offline MBO is dependent on accurate design models, and building accurate models often requires large datasets. However, collecting a large dataset with experimental measurements of PE in multiple cell types is expensive and time-consuming. In such data-constrained settings, transfer learning can be very beneficial. Indeed, Reddy et al. (2024) showed that pretraining a sequence-based model on large related genomic datasets before fine-tuning it on a smaller PE dataset from target cells leads to significantly better modelling of the smaller dataset compared to training exclusively on the smaller dataset. The three best approaches that they identified are:

### 1. Linear probing of Enformer (Avsec et al., 2021) predictions using Lasso (Tibshirani, 1996)

Enformer is a large sequence-based model (again, consisting of convolutional and transformer layers) trained on thousands of epigenomic datasets from humans and mice, allowing it to accurately learn sequence features that control gene expression. Since it is expensive to train Enformer (3 days on 64 TPU v3 cores), we often need to rely on the pretrained models published by Avsec et al. (2021), limiting flexibility in terms of changing architectures or using alternate deep learning (DL) libraries. It can also be difficult to use Lasso with gradient-based optimization techniques when performing offline MBO. Therfore, this approach is only recommended when the published pretrained Enformer model can be easily incorporated into your code base without any architectural changes, and when the offline MBO algorithm you are using is not reliant on gradients (e.g. CbAS (Brookes et al., 2019)).

### 2. Fine-tuning Enformer by using its embeddings to predict PE in target cells

The performance of this approach is very similar to the previous approach. Moreover, using finetuning instead of Lasso allows us to easily employ gradient-based offline MBO methods. Thus, this approach is recommended when the pretrained Enformer model is compatible with your code base and you do not wish to make any changes to its architecture that would necessitate re-training.

### 3. Pretraining MTLucifer on large existing MPRA datasets before fine-tuning it to predict PE in target cells

Certain MPRA datasets (e.g. (Ernst et al., 2016; van Arensbergen et al., 2019)) measure PE in selected cell lines from a large number of sequences. These data are useful for transfer learning, since models can learn a broad set of features that are generally useful for predicting PE. This approach is very inexpensive (33 hours for pretraining on a single Nvidia A40 GPU), which makes it easier to experiment with architectures and hyperparameters. Hence, this approach is recommended when you want to use custom architectures for modelling your data, or if you want flexibility in the DL libraries used to build your code base.

Using any of these approaches should yield an accurate model of PE in our target cells. However, since we focus on performing offline MBO using COMs, which require differentiable design models, only the last two approaches are compatible with our workflow.

### 4.2. Performing offline MBO while mitigating adversarial designs using COMs

Once we have accurate design models, we can couple them with offline MBO algorithms to design promoters. However, during the design process, we need to address adversarial designs that might arise due to the distribution shift problem. We describe the problem below and how it can be mitigated using COMs (Trabucco et al., 2021). Then, we specify how COMs can be used to design cell-type-specific promoters.

#### Distribution shift problem in offline MBO and conservative regularization

While we can train an accurate design model *f*_*θ*_(**x**), it may still suffer from generalization failures common to supervised regression models, especially when designed sequences are far away from the training data’s distribution i.e. when there is a distribution shift. Such adversarial sequences can be easily designed by optimizers by heavily mutating training sequences, especially those that are already good. To alleviate this issue, we suggest learning and using *conservative* models of the objective function, or COMs (Trabucco et al., 2021), as design models. These models are trained using a *conservative regularizer* that penalizes high predictions on the set of unseen and potentially undesirable promoters *μ*(**x**) which appear promising under the current design model, thus preventing the optimizer from designing adversarial promoters. Concretely, conservative models use the following loss:

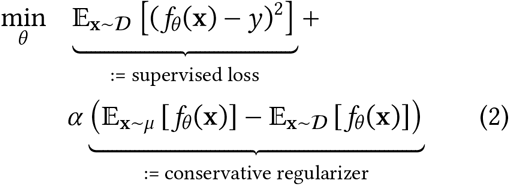

where *α* **is the conservatism coefficient** and controls the degree of “conservatism”.

#### Training design models for designing cell-type-specific promoters using conservative regularization

The previous subsection mentions two approaches for building differentiable design models. In either approach, we have a pretrained model (either a pretrained Enformer or MTLucifer model) that is finetuned using a small dataset from the target cells. To incorporate conservative regularization into the finetuning process, we modify the loss. Let us denote the model prediction for PE in cell type *c* as 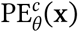, and the predicted DE in *c* as 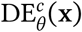, then the overall fine-tuning objective is:

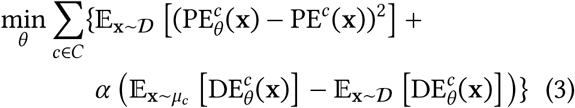

In contrast to the canonical formulation of COMs in Eqn 2, we use DE in the regularization term, a value derived from the model’s expression predictions, instead of using expression itself, as our goal is to maximize DE and we want to avoid adversarial sequences with erroneously high predicted DE using the conservative regularizer. Furthermore, since we use a single design model for all target cells, the regularization term needs to be computed for all target cells. In contrast, the canonical COMs formulation works for only one objective function. Similar to Trabucco et al. (2021), to get distribution *μ*_*c*_(**x**) consisting of sequences with potentially overestimated DE in *c* in every training step, we can use the Adam optimizer (Kingma and Ba, 2015) to perform *T* steps of gradient ascent on 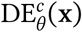 starting from sequences in the training batch (in our experiments: learning rate = 0.5, *β*_1_ = 0.9, *β*_2_ = 0.999, and *T* = 100). Since the DNA sequences are discrete, we perform gradient ascent in the probability simplex in the one-hot encoded space, parameterized by a softmax function. At the end of *T* steps of gradient ascent, we perform a hard clipping so that the resulting sequence is a valid one-hot encoding. The full fine-tuning objective is optimized till convergence and the model checkpoint with the lowest validation set loss is used as the design model. *Importantly, it is difficult to choose an α a priori. Therefore, we train multiple design models with different α values and use all of them to design candidate promoters (in our experiments, α* ∈ {0, 0.0003, 0.001, 0.003, 0.01, 0.03}).

#### Designing promoters using gradient ascent

After training a design model, starting from each promoter in the fine-tuning dataset (henceforth referred to as the **starting sequence**), we apply the same gradient ascent-based optimization process as that used to build *μ*_*c*_(**x**) to generate a design. This generation is performed using every design model (trained using different *α* values), yielding a large pool of candidate designed sequences.

### 4.3. Balancing diversity, optimality, and uncertainty for final sequence selection

From the design process described in the previous subsection, we get a large pool of candidate celltype-specific promoters. Unlike typical offline MBO problems where an algorithm must produce only a single design for evaluation, as discussed in Section 3, designed promoters are typically evaluated in batches (say, of size *K* ). Moreover, multiple iterations of promoter design can be performed. In this section we describe an algorithm to choose *K* promoters for experimental validation from the large set of candidates for cell type *c*, accounting for these facets of the promoter design problem.

Given that we have multiple design models, we cannot simply take the top *K* designs with highest predicted DE, since the predictions come from differ-ent models, making them difficult to compare with each other. Taking the top designs might also yield sequences that are very similar to each other. Finally, we also want to account for model uncertainty during selection and ideally choose high performing sequences that have low uncertainty.

### Using an ensemble to get pessimistic estimates of DE that account for model uncertainty

To uniformly evaluate all candidate designed sequences, and potentially get more accurate predictions, we build an ensemble model consisting of models with slightly different architectures, using a trainvalidation-test split of the fine-tuning data that is different from the one used to build the design models. The constituent models are still pretrained prior to fine-tuning, but the conservative regularizer is not used during fine-tuning since these models are not directly used for design. To reduce the computational overhead for building the ensemble, one can use the same pretrained backbone in all constituent models but use different kinds of fine-tuning layers in each model. In our experiments, we use 36 models in the ensemble. To account for model uncertainty, we use the ensemble to compute 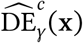, a pessimistic estimate of the DE in cell type *c*, for all candidates. The pessimistic estimate is defined as the lower confidence bound of the constituent models’ predictions and is computed as the mean minus the standard deviation of the predictions. *Using this pessimistic estimate in the final selection process allows us to choose sequences that most models are confident about, further reducing the risk of choosing adversarial designs*.

### Final selection algorithm

Our ultimate goal is to maximize the expected efficacy of the designs in the set of final sequences selected for experimental validation. Intuitively, this can be achieved by: **(i)** ensuring that our designs have high predicted DE, and **(ii)** ensuring that the designs are as diverse as possible. Diversity is important as designs predicted to be optimal might not actually be optimal when they are experimentally validated. Having a diverse set of sequences increases the likelihood of *some* design being optimal, since we cover a broad region of the sequence space. We define a sequence set 𝒮’s diversity as:

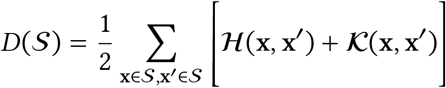

where ℋ is the normalized Hamming distance between two sequences (ensuring overall diversity) and 𝒦 is the normalized Euclidean distance between the 6-mer frequency vectors of two sequences (ensuring diversity in potential TF-binding motifs composition). The distances are normalized by dividing them by their maximum possible values. To instantiate this intuition into a concrete strategy, we aim to select a set of *K* sequences 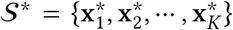 such that 𝒮^*^ is the optimal solution to the following optimization problem:

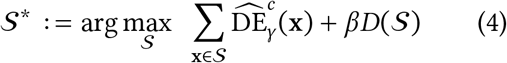

where 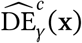 denotes a pessimistic estimate of the DE computed using an ensemble, and *β* **is the diversity coefficient** that controls the fitness vs. diversity trade-off. While in theory we can obtain 𝒮 ^*^ by optimizing over all sequences, doing so is computationally intractable. So, the optimization in Eqn 4 is performed using a greedy algorithm over a large subset of candidates from the previous step. The subset consists of candidates that have positive ensemble average predicted DE, and also have high predicted expression in the target cell type *c* ^1^ since this is also crucial for gene therapy applications. The algorithm sequentially chooses *K* sequences - in each step, given a set of sequences 𝒮^′^ already chosen to be in the final set, it computes 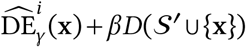 for every candidate sequence **x** not in 𝒮^′^, and chooses the sequence that maximizes this value for inclusion in the final set.

### 4.4. Making adjustments to the design process based on performance and diversity metrics

Here we discuss common problems that may arise during design and provide heuristics to address them.

#### Designed sequences are generally not predicted to have higher DE than the starting sequences from the fine-tuning dataset

This issue might arise when the conservative regularizer is too powerful, making it difficult for the optimizer to design good sequences. Here, reducing the conservatism coefficient *α* should help. However, there might also be problems with the dataset that hinder the optimizer. For example, if the fine-tuning dataset has very few sequences that drive some level of DE in the target cell type, the design models might be poor at modelling such sequences. To determine if this is the case, one can look at the distribution of DE values in the training set and if there are too few sequences that have positive DE in the target cell type (e.g. less than 25% of the dataset), the design problem might be too difficult to solve using the available data.

#### Designed sequences are not diverse

The diversity of a set of designs can be measured using multiple metrics. We focus on base pair entropy and average pairwise edit distance (can be any edit distance e.g. Hamming distance) as they are easy to interpret. If we are designing sequences of length *L*, base pair entropy can be computed at every position in the sequence by determining the frequency of the 4 DNA bases across all designed sequences. For a diverse sequence set, this metric should be close to 2 at every position (near uniform usage of the 4 bases). The average pairwise edit distance should be comparable to *L* (e.g. *L*/2). If these metrics are low, the diversity coefficient *β* can be increased to boost diversity. Using a more diverse set of starting sequences during optimization with gradient ascent can also improve the diversity of the designed sequences since we can converge upon different optima of the objective landscape.

## 5. Implementation and experimental evaluation of our approach

Next, we use the workflow proposed in the previous section to design 250 bp long cell-type-specific promoters for three relatively similar blood cancer cell lines: Jurkat, K562, and THP1. As mentioned in the introduction, designing such promoters is difficult since these cell lines are very similar compared to cell lines derived from different tissues. Moreover, Jurkat and THP1 cells are relatively understudied and lack large sources of PE measurements. We also perform experimental validation of the designed sequences to show the effectiveness of our approach.

### 5.1. Setup

#### Model training (design and ensemble models)

To train design models, we use an architecture similar to MTLucifer (Reddy et al., 2024) (shown in Figure S.1). We pretrain the models on large existing MPRA datasets before fine-tuning them using a small PE dataset consisting of 17,104 measurements collected by Reddy et al. (2024) from Jurkat, K562, and THP1 cells (dataset is described in more detail in Appendix B). We use this approach over fine-tuning Enformer, since our codebase uses Jax (Bradbury et al., 2018) for efficiently performing conservative regularization, and pretrained Enformer models are not available for Jax. Following Section 4.2, we train 5 design models, each with a different *α* value. We also build an ensemble for final sequence selection using a different split of the same dataset. Appendix D describes the training process in more detail.

#### Sequence design

We design 4,000 sequences for each cell type using our workflow. This number was chosen based on experimental constraints, and to keep the total number of sequences close to the size of the original training dataset. Each design model is used to design *∼*17K candidate sequences (one design for every starting sequence from the fine-tuning dataset) using the process described in Section 4.2, and the final 4,000 sequences are chosen from these candidates using the algorithm from Section 4.3. As shown later in this section, the designed sequences are naturally diverse, and we set the diversity coefficient *β* to zero since we did not need to improve diversity.

#### Baselines

We also design sequences using two baselines, in order to determine the usefulness of various components of our workflow:

##### 1. Motif tiling

This heuristic method to design cell type-specific promoters is similar in spirit to that proposed by Nissim et al. (2017), and aims to improve upon known high DE sequences. We use it to determine the overall usefulness of our approach for designing promoters compared to traditional methods. Briefly, the motif tiling approach inserts motifs associated with high DE into sequences from the fine-tuning dataset that already have high DE, with the goal of increasing their DE even further. We run it for the three target cell types to design 520, 630, and 515 sequences for Jurkat, K562, and THP1 cells, respectively. Appendix E.1 presents the details of this method.

##### 2. DENs

In Section 4.2, we use gradient ascent to optimize starting sequences and produce designs that increase the design model-predicted DE. However, as detailed in the Section 2, many previous studies use generative models for this purpose. To determine which optimization algorithm is better, we replace gradient ascent with DENs in our workflow and produce 2000 sequences per cell type (with *β* = 10). We specifically use DENs, as their designs were experimentally validated by Linder et al. (2020) and they are explicitly trained to produce diverse sequences. Appendix E.2 describes DENs in more detail.

#### Experimental validation of sequences

We use a similar protocol as Reddy et al. (2024) to experimentally determine the DE of designed sequences (Appendix C). Along with the designed sequences, we also measure the DE of the top sequences from the training dataset (top 100 sequences for each cell type, so 300 sequences total), and of *∼*200 sequences from the training dataset whose expression roughly uniformly spanned the full range of PE values. These sequences with measurements in both the training set experiment and the validation experiment are used to determine whether designed sequences improve upon their starting sequences, using a procedure described later in this section. In total, we measure the DE of 20,741 sequences. After filtering out sequences that have less than 100 reads in any experiment, we are left with 14,315 high-quality measurements for use in all downstream analyses.

### 5.2. Main Results

#### Evaluating the diversity of designed sequences

We analyze the diversity of the designs in terms of their mean base pair entropy and the mean Hamming distance between any two designs (Table 1). **Overall, our approach produces highly diverse sequences for all three cell types**. We note that for THP1, DENs produce significantly less diverse designs. This is possibly due to the difficulty of the design problem - the fine-tuning dataset has few sequences with high DE in THP1, leading to the design model having fewer modes. This might make it difficult for a generative model trained from scratch to discover the design model’s modes. Since gradient ascent starts from the sequences in the finetuning dataset, it can more easily discover the various modes.

**Table 1:**
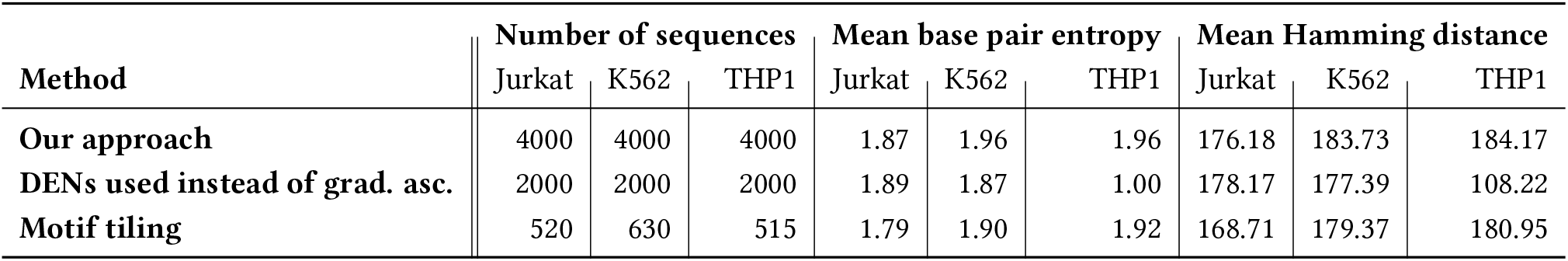
Quantifying the diversity of designs. Mean base pair entropy (maximum value = 2) and mean pairwise Hamming distance (maximum value = 250) are shown for the designed sequences. Mean base pair entropy is calculated by determining the entropy of each position across all designs and then averaging these values.

#### Does our approach design sequences that are better than the corresponding starting sequences?

Next, we analyze the overall effectiveness of our workflow and compare it to motif tiling. Since we aim to design promoters that have higher cell-type-specificity than promoters in the fine-tuning dataset, we need to quantify the improvement in DE from the starting sequences to the designed sequences. Direct comparison of DE values is challenging due to differences in experimental conditions and potential confounding factors like batch effects, and re-measuring the DE of all starting sequences is inefficient. To address this issue, we use a common set of sequences that span the expression range for any given cell type to calibrate DE values between experiments. Since these common set sequences have DE values from both experiments, we can compute percentile scores for both the starting and corresponding designed sequences among this common set. The design process is considered effective if the designed sequence achieves a higher percentile score than the starting sequence. This calibration is valid since the ranks of the sequences in the common set are concordant between experiments, as evidenced by the high Spearman correlation between DE values measured by Reddy et al. (2024) and our experiments: 0.953 for Jurkat, 0.921 for K562, and 0.950 for THP1.

Figure 2 shows the percentile scores of the starting and designed sequences from our approach and from motif tiling. Sequences designed for Jurkat and K562 using our approach improve upon the majority of corresponding starting sequences, highlighting the overall effectiveness of our approach in this difficult setting. **In fact, we design a promoter for K562 that has 75.85% higher DE (=5.17) than the best high DE sequence from the fine-tuning dataset (=2.94)**. On the other hand, we observe that using motif tiling to improve on high DE sequences is ineffective, almost always leading to worse sequences. We note that our workflow tends to choose designs that come from relatively mediocre starting sequences. This is possibly because most sequences in the fine-tuning dataset have DE similar to these sequences, likely making the design and ensemble models accurate and confident near such sequences. Since our workflow significantly improves upon such sequences, we should be able to discover progressively better sequences over multiple rounds of model training, design, and experimental evaluation.

**Figure 2:**
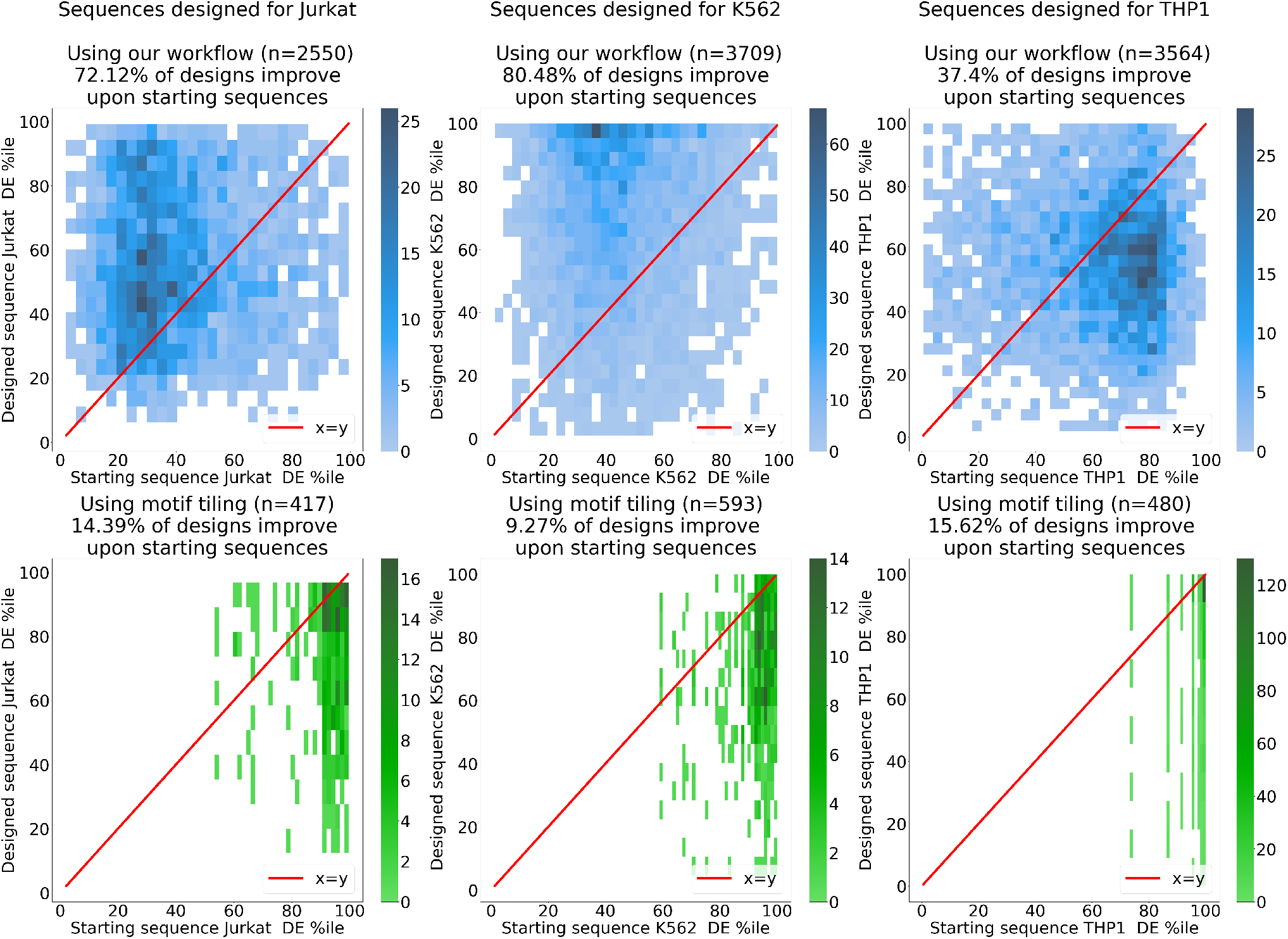
Comparing the DE of designed sequences with the DE of corresponding starting sequences. The top row of plots compare the percentile scores of the starting and designed sequences from our approach, and the bottom row of plots compare the percentile scores for starting and designed sequences from motif tiling. Each column represents sequences designed for one of the three cell types.

However, we fail to improve upon most starting sequences for THP1. This is likely because of the difficulty of the design problem. As mentioned previously, there are few high DE sequences in the finetuning dataset for THP1. Thus, it might be difficult for design models to be accurate in the space of high DE sequences, leading to poor designs. In such cases, collecting more experimental data for training might be the only effective way to improve designs.

#### Gradient ascent is better than DENs for designing sequences

Finally, we compare gradient ascent to DENs (Figure 3) to determine the better optimizer for use in our workflow. We observe that gradient ascent is generally more effective than DENs. Given that gradient ascent is also easier to implement and tune, we recommend it over generative models such as DENs.

**Figure 3:**
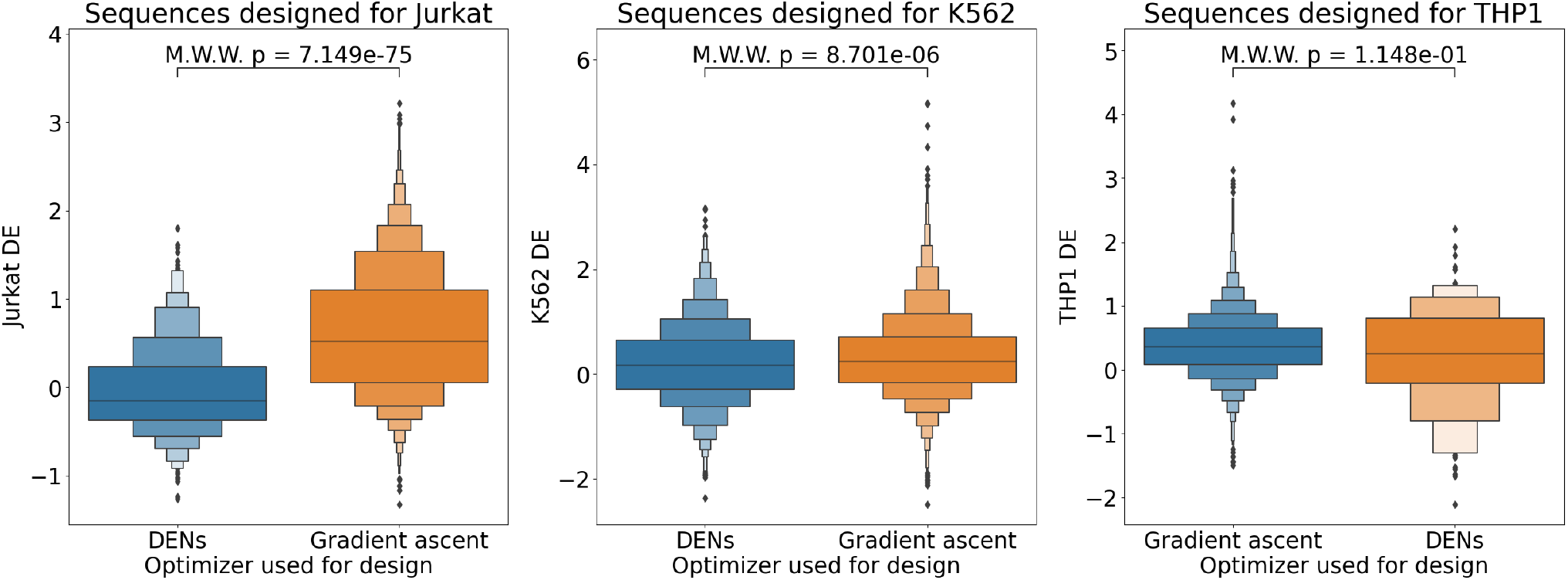
Box plots comparing the DE of sequences designed using gradient ascent as the optimizer in our workflow vs. those from DENs. *p*-values are computed using a Mann-Whitney-Wilcoxon (MWW) test that tests whether sequences from gradient ascent have higher DE than sequences from DENs.

## 6. Discussion and limitations

Designing cell-type-specific promoters is a crucial step in effective application of gene delivery technologies. We introduce a novel workflow for promoter design based on COMs (Trabucco et al., 2021), tailored for real-world applications due to its data efficiency and ability to produce diverse, non-adversarial designs. We apply this approach to the challenging task of designing cell-type-specific promoters for three similar blood cancer cell lines. We then experimentally validate our designs, demonstrating effectiveness for two out of three cell lines.

We also present limitations that practitioners should consider. The success of our workflow depends on the accuracy of the design models, and effectiveness across the three cell lines was related to design model accuracy for each cell line (Table S.1). This resulted in failure for THP1 cells, where we had lower quality fine-tuning data with few high DE sequences. Pretraining data may also affect performance, and many existing MPRA datasets used for pretraining come from K562 cells, likely explaining our superior performance in K562 compared to Jurkat. Our MBO workflow highlights these and other practical considerations. Our approach is valuable for practitioners targeting a wide range of cell types, and will inspire further research in this area.

## Acknowledgments

This work was partially supported by an H2H8 Graduate Research Grant to AJR, a Schmidt Futures grant to SL, and the U.S. National Institutes of Health grant R00HG009677 to NMI.

This research used the Savio computational cluster resource provided by the Berkeley Research Computing program at the University of California, Berkeley (supported by the UC Berkeley Chancellor, Vice Chancellor for Research, and Chief Information Officer).

## A. Supplementary Figures

**Figure S.1:**
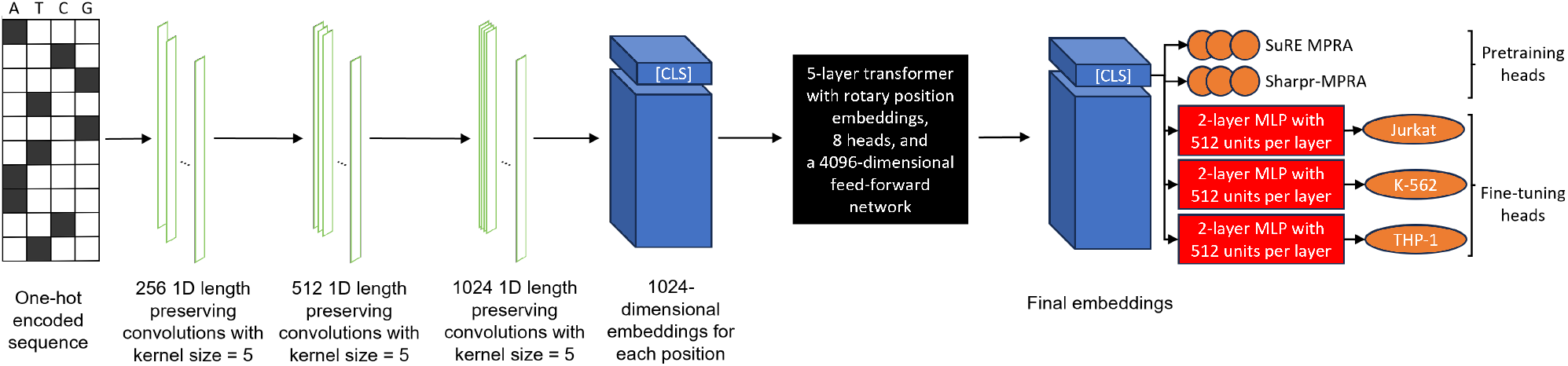
Design model architecture: the convolutional layers use GELU activation (Hendrycks and Gimpel, 2016), dropout with 0.1 probability (Srivastava et al., 2014), and are followed by a group norm layer (Wu and He, 2018) with each group size being 16. The MLP layers in the fine-tuning heads also use GELU activation. We apply the RoPE position embeddings (Su et al., 2021) at each attention layer of the transformer.

## B. Experimental data used for training design models

We use the PE measurements collected by Reddy et al. (2024) for designing cell type-specific promoters. They provide 17,104 PE values in Jurkat, K562, and THP1 cells derived from 250 bp long manually designed promoters. They set out to measure the PE of 20,000 promoters chosen using heuristics that try to maximize the number of differentially expressed promoters. Nearly 50% of the tested promoters are from differentially expressed endogenous genes, another *∼* 40% are crafted by assembling known and de-novo motifs which are abundant in the promoters of differentially expressed endogenous genes, and the remaining *∼* 10% of promoters originate from highly expressed endogenous genes’ promoters. Each promoter is integrated upstream of a minimal CMV promoter and the enhanced green fluorescent protein (EGFP) reporter gene within a lentiviral vector. The resulting expression in each cell line post-transduction is quantified by the levels of induced fluorescence. They get 17,104 PE values of adequate quality with two replicates and average the values from each replicate to get the final PE values - we use these values in our experiments.

This data was released with their codebase (https://github.com/anikethjr/promoter_models) under an AGPL-3.0 license.

## C. Experimental protocol for measuring promoter-driven expression of designed sequences

We used the same protocol as Reddy et al. (2024) with the following modifications for testing the newly designed sequences. First, the cycle number for PCR amplification of the Twist oligopool was reduced from 12 to 10 cycles to reduce amplification bias, and the lentiviral backbone used a CMV promoter instead of an RSV promoter to drive the expression of the cargo from the lentiviral transfer plasmid to increase titer. To generate lentivirus, we used the LV-MAX kit (ThermoFisher A35684) following the protocol for a 1L flask, followed by the same concentration method. For lentiviral titration and full scale transduction, we used the same protocol, except we did not perform spinfection as this was deemed not necessary for adequate transduction efficiency. After transduction, medium containing 8ug/mL polybrene and lentivirus was changed to fresh medium after 24 hours, and cells were allowed to recover for another 48 hours before performing puromycin selection. This additional time allowed for better expression of the puromycin resistance gene before selection, increasing functional titer. Full scale transductions were performed at >500X library coverage and at an MOI <0.3. Cells were sorted using the same general methodology at a similar coverage; however, gDNA was extracted using the NucleoSpin Blood L (Machery-Nagel 740954) with the addition of RNAse A (NEB T3018L) following the manufacturer’s instructions. No changes were made to NGS library preparation, but sequencing was performed using an Illumina NovaSeq X Series, pooled with other NGS libraries with non overlapping indices.

## D. Training of design and ensemble models and their prediction performance

In this section, we detail the process we use to train our design and ensemble models. We train 5 design models with varying values of the conservatism coefficient *α* (*α* ∈ {0, 0.0003, 0.001, 0.003, 0.01, 0.03}). We also highlight their prediction performances.

### D.1. Pretraining details

Here, we provide details about our pretraining process. All design models and all constituent models of the ensembles are pretrained before being fine-tuned. Following Reddy et al. (2024), we pretrain our models using data from Sharpr-MPRA (Ernst et al., 2016) and SuRE MPRA (van Arensbergen et al., 2017; 2019). SuRE MPRA measures the expression induced by 150-500bp genomic fragments from 4 individuals from 4 different populations in the K562 and HepG2 cell lines. *∼* 2.4B and *∼* 1.2B fragments were found to be expressed in K562 and HepG2 respectively. Most fragments have very low expression and training on all measurements is time-consuming. Thus, Reddy et al. (2024) define a classification task using this data that subsets the data and bins each sequence into one of 5 expression bins. We also use this classification task for pretraining. Sharpr-MPRA is a smaller MPRA that measures the expression from *∼* 487K 145bp sequences centered at DNase I peaks in K562 and HepG2 cells and in two different settings. Reddy et al. (2024) use a preprocessed version of this data from Movva et al. (2019) that builds a regression task with 12 outputs (2 replicates for expression measured in 2 settings in 2 cell lines, and 4 outputs that correspond to the average expression across replicates). We use the same formulation for pretraining our models.

During pretraining, our models’ output layers produce the probability of a sequence belonging to each of the expression bins for the SuRE MPRA-based pretraining task and directly predicts the expression values for the Sharpr-MPRA-based pretraining task. We minimize the sum of the negative log-likelihood (NLL) loss for the SuRE MPRA task and the mean squared error (MSE) loss for the Sharpr-MPRA task (since both tasks have distinct sequences, a training sequence only contributes to one of the two loss terms, the other loss is set to zero for that sequence).

We use the dataset splits defined by Reddy et al. (2024) and train with a batch size of 448 using the AdamW optimizer (Loshchilov and Hutter, 2017) with 1e-4 learning rate and 3e-3 weight decay. We train the models for 20000 steps in total, with a cosine learning rate decay schedule that decays to 0. We retain the best checkpoint that is selected by monitoring the validation loss throughput the training process.

### D.2. Building an ensemble by combining models with slightly different architectures

The ensemble uses constituent models that slightly modify the design model architecture presented in Figure S.1 - each model uses a different type of MLP layer (shown in red in Figure S.1) in the fine-tuning head. Within the set of constituent models, we vary the depth (2, 4, or 8 layers), number of hidden units (512, 1024, or 2048), and activation functions (tanh, GELU, ReLU, or SiLU (Hendrycks and Gimpel, 2016)) of the MLP layers. This gives us 36 constituent models for ensembling.

### D.3. Fine-tuning details

After pretraining, all design models are fine-tuned using a certain split of the PE dataset collected by Reddy et al. (2024) (Split 1 in Table S.1). The ensemble’s constituent models are fine-tuned using a different split (Splits 2 in Table S.1). In each split, following Reddy et al. (2024), we use *∼* 70% of the assayed promoters for training, *∼* 10% for validation, and *∼* 20% for testing. Since there are distinct classes of promoters in the dataset with varying levels of GC content, our splits are stratified by both promoter class and GC content.

When fine-tuning the design models using the conservative regularizer, the full fine-tuning objective from Eqn 3 is optimized for up to 200 steps using the AdamW optimizer (Loshchilov and Hutter, 2017) (batch size = 512, learning rate = 5e-5, weight decay = 3e-3, *β*_1_ = 0.9, *β*_2_ = 0.999) with a cosine learning rate decay schedule that decays to 0 after 10 warm-up steps and model checkpoints with the lowest validation set loss are retained.

The constituent models of the ensemble are fine-tuned without the conservative regularizer using the AdamW optimizer with a batch size of 512, 5e-5 learning rate, 3e-3 weight decay, *β*_1_ = 0.9, *β*_2_ = 0.999, and with a cosine learning rate decay schedule that decays to 0 after 10 warm-up steps. They are fine-tuned for upto 250 steps, and we retain the best checkpoints that are selected by monitoring the validation loss throughput the training process.

### D.4. Prediction performance

The prediction performance of our models on their respective test sets is shown in Table S.1. These results indicate that all of our models are accurate predictors of PE in the target cell types, justifying their usage in designing promoters.

**Table S.1:** Prediction performance obtained using our models.

| Model | Dataset Split | Jurkat |  | K562 |  | THP1 |  |
| --- | --- | --- | --- | --- | --- | --- | --- |
| | | $r$ | $\rho$ | $r$ | $\rho$ | $r$ | $\rho$ |
| Design model with $\alpha = 0$ | Split 1 | 0.699 | 0.666 | 0.705 | 0.698 | 0.644 | 0.542 |
| Design model with $\alpha = 0.0003$ | Split 1 | 0.700 | 0.666 | 0.706 | 0.698 | 0.644 | 0.542 |
| Design model with $\alpha = 0.001$ | Split 1 | 0.700 | 0.666 | 0.706 | 0.698 | 0.643 | 0.542 |
| Design model with $\alpha = 0.003$ | Split 1 | 0.701 | 0.667 | 0.706 | 0.698 | 0.642 | 0.543 |
| Design model with $\alpha = 0.01$ | Split 1 | 0.700 | 0.668 | 0.705 | 0.698 | 0.639 | 0.545 |
| Ensemble model used for final selection | Split 2 | 0.702 | 0.659 | 0.691 | 0.690 | 0.627 | 0.529 |
| Test Set Replicate Concordance | Split 1 | 0.819 | 0.735 | 0.737 | 0.666 | 0.730 | 0.483 |

## E. Details about baselines

In this section, we describe the baselines in more detail.

### E.1. Designing cell type-specific promoters using motif tiling

In this section, we detail how we design cell type-specific promoters using motif tiling. First, to identify motifs that might contribute to DE, we use FIMO (Grant et al., 2011) with default settings to detect instances of clustered TF-binding motifs defined by Vierstra et al. (2020) ^2^ in the sequences assayed by Reddy et al. (2024), and retain detected motif occurrences with q-value < 0.01. Let’s now consider designing a cell type-specific promoter for Jurkat. For every motif, we first run a Welch’s t-test to determine if sequences that contain it have higher expression in Jurkat than sequences that do not contain it, and retain motifs with q-values < 0.01. Then, for every motif, we run 2 pairwise Welch’s t-tests to determine if its presence leads to higher expression in Jurkat compared to K562 or THP1. Motifs that have positive effect sizes in both t-tests (i.e. leads to higher expression in Jurkat compared to both K562 and THP1) with q-values < 0.01 are retained as those that could be contributing towards DE in Jurkat. Then, these motifs are used to design two sets of sequences - one set of sequences is designed by inserting the same motif into a background sequence as many times as possible while separating the motifs by 10bp (we get 5 sequences per motif using different background sequences), and the second set is designed by randomly sampling motifs (weighted by average effect size from the two pairwise tests) from the list of retained motifs and inserting as many of them as possible into a background sequence while separating the motifs by 10bp. While generating each sequence, the background sequence is a randomly chosen sequence from those assayed by Reddy et al. (2024) that exhibits a DE of at least 2 and also has PE in the target cell that is greater than the 90th percentile of PE. Inserted motif sequences are sampled from the motif’s position weight matrix (PWM). The same process is repeated for K562 and THP1.

We discover 4, 26, and 3 motifs that may be causing DE in Jurkat, K562 and THP1 respectively. Thus, we get 20, 130, and 15 sequences in Jurkat, K562, and THP1 respectively by tiling the same motif repeatedly. Then, for each cell line, we design 500 sequences by randomly sampling motifs. Therefore, we get a total of 520, 630, and 515 sequences for Jurkat, K562, and THP1 respectively using this design method.

### E.2. Deep Exploration Networks (DENs)

DENs are generative models that are trained to output diverse sequences that maximize a design model’s predictions. The generator takes random noise as input and transforms it into a sequence PWM. We use a UNet-style (Ronneberger et al., 2015) generator that first transforms the noise vector into a sequence PWM-sized matrix (i.e. of size (250, 4)). Then, it applies 6 downsizing convolutional layers followed by 5 upsizing convolutional layers. A final convolutional layer then pools information across the final set of filters’ outputs to produce the sequence PWM. The PWM is used to sample sequences that are fed to the design model to get its predictions and a fitness-based loss that trains the DEN to output high-fitness sequences is computed. Additionally, to explicitly increase the diversity of the generated sequences, in every training step, random noise vectors are input to the DEN in pairs to get two sequence PWMs per pair. Then, a diversity-based loss is computed that incentivizes the sequences generated using the two different noise vectors to be distinct from each other, both in sequence and design model embedding space. An entropy-based loss is also minimized to reduce the entropy of the PWM output by the DEN at every position.

Thus, when training a DEN to generate cell type-specific promoters for a target cell *i* ∈ {Jurkat, K562, THP1}, the training objective we use is:

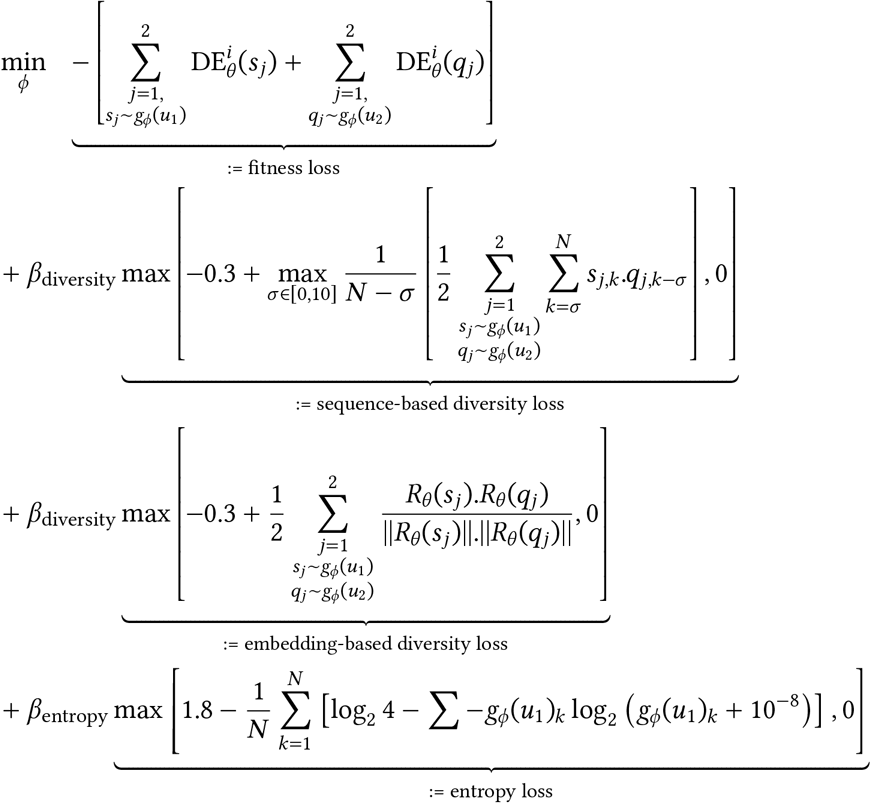

where *ϕ* is the set of trainable parameters of DEN *g*_*ϕ*_ which outputs *N* = 250 base pairs long sequence PWMs by taking *u*_1_ or *u*_2_ - 200-dimensional random noise vectors sampled from the uniform distribution over [-1, 1], as inputs. From the sequence PWMs output by *g*_*ϕ*_, we sample two one-hot encoded sequences per noise vector denoted by *s*_*j*_ and *q*_*j*_ . These sequences are then input to the trained design model that predicts PE induced in each of the three cell types. Then, the predicted DE in the target cell *i* induced by a sequence **x** is given by Eqn 1. The fitness loss maximizes this predicted DE. The other loss terms increase sequence diversity and reduce entropy in the sequence PWM. Here, *s*_*j*,*k*_ is the one-hot encoded base pair at position *k* in *s*_*j*_ (similarly for *q*_*j*,*k*_), *R*_*θ*_(*s*_*j*_ ) is an embedding for *s*_*j*_ extracted from the design model *f*_*θ*_, *g*_*ϕ*_(*u*_1_)_*k*_ is the probability distribution over base pairs in the sequence PWM *g*_*ϕ*_(*u*_1_) at position *k*. Finally, the coefficients *β*_diversity_ and *β*_entropy_ are used to weight the diversity and entropy losses relative to the fitness loss and to one another. They can be varied to regulate the diversity vs. fitness trade-off. We refer readers to the original work by Linder et al. (2020) that proposed DENs for more details on the method. We tune the various hyperparameters reflected in the training objective by observing the overall quality of the generated sequences.

We train a total of 15 DENs per target cell type, each using a different design model to compute the fitness loss, or making a different diversity vs. fitness trade-off by using different *β*_diversity_ values. We have 5 different design models, each trained using a different conservatism coefficient *α* (Table S.1). We also try 3 different *β*_diversity_ values - 1, 5, and 10, yielding 15 DENs in total. Each DEN is used to generate 20000 sequences and we use the final sequence selection algorithm from Section 4.3 to choose the final set of 2000 sequences per target cell.

## Footnotes

1 Designs with ensemble average predicted PE in target cell type *c* greater than the 90th percentile of predicted PE in *c* for the fine-tuning dataset’s sequences.

2 https://resources.altius.org/~jvierstra/projects/motif-clustering-v2.0beta/

